# Altering subjective time perception leads to correlated changes in neural activity and delay discounting impulsivity

**DOI:** 10.1101/2024.01.14.575587

**Authors:** Sangil Lee, Joseph W. Kable, Wi Hoon Jung

## Abstract

Several accounts of delay discounting suggest subjective time perception as a contributing factor to individually varying discount rates. That is, one may seem impatient if their subjective perception of delay is longer than others’ perception of it. Here we build upon the behavioral and neural research on time perception, and we investigate the effects of manipulating an individual’s subjective time perception on their discount rates and neural activity. Using a novel time-counting task, we found that participants’ discount rates are affected by our manipulations of time perception and that neural activity also correlates with our manipulations in brain regions, such as the anterior insula and the superior temporal gyri, which have been implicated in time perception. We link these behavioral and neural findings together by showing that the degree of neural activity change in response to our manipulation is predictive of the degree of change in the participants’ discount rates.

---

Time, though it may be but a stubborn illusion, is an inextricable part of human decision-making. It permeates our daily lives as we decide whether to expedite packages for extra cost, to eat healthy and quit smoking for future health, or to budget for immediate spending to enable saving for decades-long retirement plans. In all these decisions, we humans are known to vary considerably in how much we devalue future rewards compared to immediate ones – a phenomenon known as delay discounting. The rate at which we devalue future rewards (i.e., the discount rate) has been shown to predict a growing list of real-life metrics such as school grades, education level, smoking, substance abuse, and financial status (Alessi & Petry, 2003; Atlas et al., 2017; Kirby et al., 1999, 2005; Schepis et al., 2011; Shamosh & Gray, 2008; Urminsky & Zauberman, 2015).

So why are some of us more impatient than others? The breadth of the question invites quite a number of theories, but perhaps one of the simplest is the account of subjective time perception: we don’t perceive time in the same way. (Bilgin & LeBoeuf, 2010; Ebert & Prelec, 2007; B. K. Kim & Zauberman, 2019; K. Kim & Zauberman, 2019; Zauberman et al., 2009). If one’s subjective perception of time is slower than others, the same delay would feel longer and would result in a higher discount rate in objective time. The subjective time perception theory also has the benefit of being able to rationally account for the hyperbolicity of the discounting function by combining the economically normative exponential function with non-linear time perception (Cui, 2011; Ray & Bossaerts, 2011; Zauberman et al., 2009).

Despite the intuitiveness of the theory of subjective time perception, the neural link between time perception and discount rate has not yet been investigated. On the behavioral side, previous research has used within-subject manipulations such as time pressure (Ebert & Prelec, 2007) or music tempo (K. Kim & Zauberman, 2019) to influence peoples’ perception of time and, as a consequence, their discount rates. However, the neural effects of these time perception manipulations have been less well studied. What has been studied on the neural side, using a duration-reproduction task, are neural ‘pace-keeping’ regions such as the insular cortex (Bueti & Macaluso, 2011; Wencil et al., 2010; Wittmann et al., 2010, 2011). Several studies have shown that those with higher neural activity in the insular regions during a duration-reproduction task generally reproduced shorter intervals than those with lower activity (Bueti & Macaluso, 2011; Wittmann et al., 2011). However, these neural findings were established across subjects as a trait-like measure of time perception have yet to be linked to within-subject manipulations in time perception or to behavioral changes in delay discounting.

In this study, we seek to bridge the gap in the literature by experimentally manipulating time perception and measuring both behavioral changes in delay discounting and neural changes in activity. We provide a novel ‘time-counting’ task that builds upon the duration-reproduction task to directly manipulate the pace at which people count time. The time-counting task is intermixed with an intertemporal choice (ITC) task to dynamically measure discount rates over time. First, we show that our manipulation in the time-counting task changes participants’ discount rates. Second, we show that the same manipulation also causes changes in neural activity in multiple regions, some of which have been previously implicated in time-keeping. Finally, we show that the degree of neural activity change in these brain regions predict the degree of change in discount rate.

## Methods

### Participants

Thirty-eight people participated in the experiment with informed consent. Two requested early termination of the study due to nausea. One participant repeatedly did not respond to the task during the scan and was excluded for non-compliance. Three additional participants who completed the tasks were excluded from data analysis as their choices in the ITC task were extremely one-sided, prohibiting us from reliably estimating their discount rates (see below for details). This resulted in a total of 32 final participants for this study (20 males, 12 females; 19- 29 years old, mean = 21.84 yrs, SD = 2.70 yrs). All experimental procedures were approved by the Institutional Review Board of Daegu University.

### Experimental Design

To measure the relationship between time-counting and delay discounting, we alternated between time-counting and ITC tasks such that one trial of a time-counting task was followed by 3 trials of an ITC task. In the time-counting task, participants saw a series of numbers appearing on the screen sequentially and rhythmically from 1 to 5, after which the screen went blank but for a fixation cross in the center. Participants were asked to keep track of the tempo at which the numbers increased from 1 to 5 and to push the button on their button box at the precise moment they felt like the number would have reached 12 (**Figure 1A**). To manipulate the participants’ internal time-counting tempo, we varied the pace at which the numbers counted from 1 to 5 across three conditions: in the normal condition, the numbers increased every second, while in the fast and slow conditions the numbers increased every 0.7 and 1.3 seconds, respectively. Hence, in the normal condition, participants should press the button 7 seconds after the number 5 appears, while in the fast condition, 4.9 seconds, and in the slow condition, 9.1 seconds. The response screen lasted for a little longer than the correct timing to allow for measuring late responses: 10.2s, 12s, and 14.8s in fast, normal, and slow conditions.

**Figure 1.**
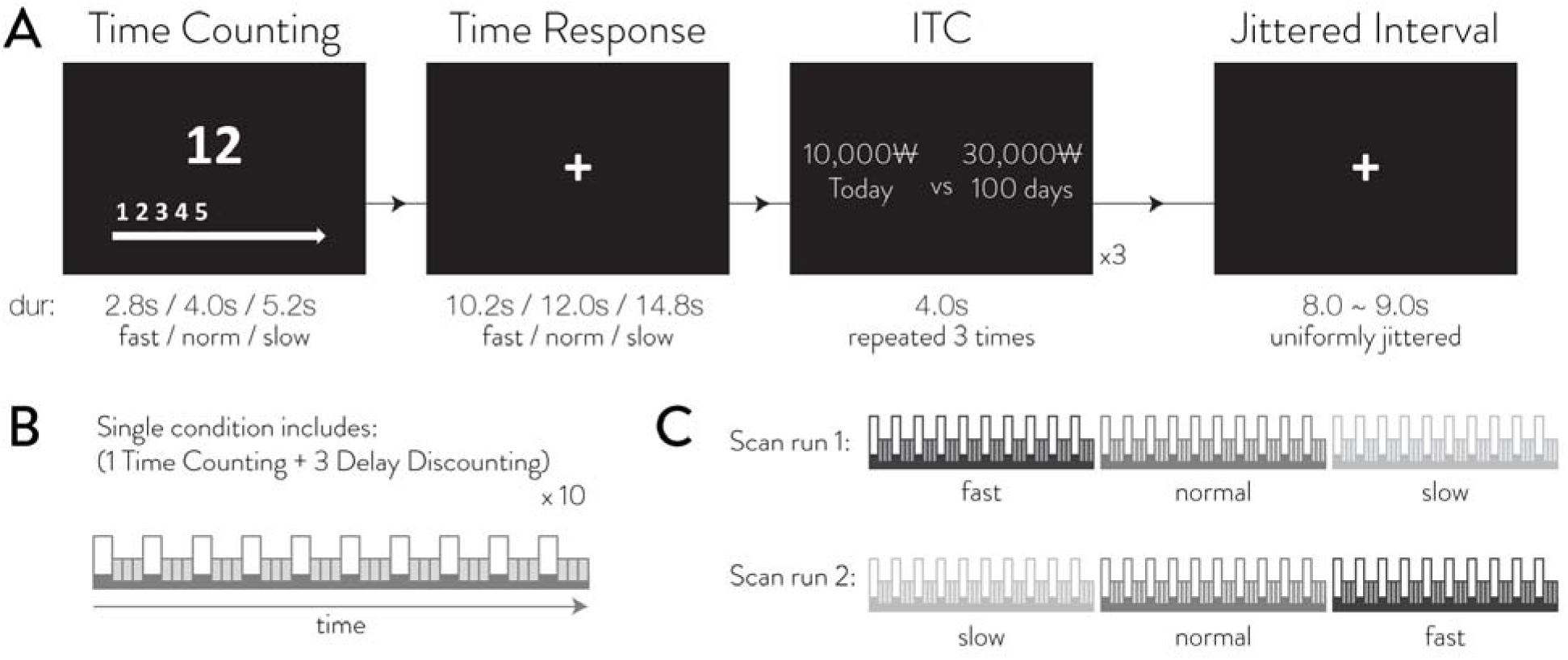
Task description. Panel A shows the task screens of a single experimental block, which consists of one time-counting task trial and three intertemporal choice (ITC) task trials. Panel B depicts a single experimental condition that consists of 10 experimental blocks like the one shown in panel A. Panel C shows the condition layout of the two fMRI scan runs, with the first run involving fast, normal, and slow conditions, and the second run in reverse order.

Following one trial of time-counting task, participants completed three trials of a binary ITC task in which they made decisions between a smaller immediate monetary reward of 10,000 Korean won (roughly $8∼$9 USD) and a larger delayed monetary reward that varied in amount and delay from trial to trial (**Figure 1A**). The larger delayed amount varied from 11,000 won to 60,000 won and the delay varied from 2 days to 180 days. The delays and the amounts were sampled such that we get a uniform sampling of the log of the hyperbolic discount rate: *k* (Green & Myerson, 2004; Mazur, 1987). Given a delayed amount *A* and delay, the hyperbolic discounting function gives us the delayed utility of *A/(1+k*σ*i*, which if equal to 10,000 won, would mean that the discount rate *k* would equal *1(A/10000i-1 /*σ**. We varied the amount and delay such that *,nk* (natural log of *k*) would be sampled uniformly in the range of -7.5 and 1 across 60 trials. These 60 trials were repeated for each of the three conditions.

All participants completed two fMRI scan runs. Each scan run consisted of three conditions (**Figure 1C**), each with 40 trials (10 time-counting trials + 30 delay discounting trials; **Figure 1B**). The first scan run’s conditions were in the order of fast, normal, and slow and the second scan run’s conditions were in the reverse order of slow, normal, and fast.

### Behavioral Data Analysis

We estimated participants’ discount rate by fitting a Bayesian hierarchical binary logit model of the following form. The subscript *i* denotes the individual, and, denotes the trial.

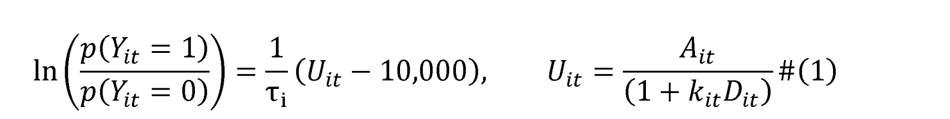

**Eq. 1** shows that the log odds of individual *i*; choosing the delayed reward on trial, (*Y*_*i*_ *1*) is modeled by the difference in utility between the delayed option (*u*_it_) and the immediate option, whose utility is fixed at 10,000 won. The *T*_i_ is the individual softmax inverse temperature parameter that controls the noisiness of the behavior (infinite temperature resulting in stochastic 50/50 choices and 0 temperature converging to deterministic choice based on utility). The utility *U_it_* is modeled by a hyperbolic discounting function where the delayed amount of the trial, *A*_it_, is divided by *1+k*_it_*σ*_it_ where *σ*_it_ is the delay of the trial and *k*_it_ is the discount rate of the trial.

The first goal of this experiment is to identify how the time perception manipulation affects the discount rate. We considered four different possibilities, not all of which are time perception related, by which our time-counting conditions could affect the discount rates. These were used as regressors to model the change of log discount rate over time:

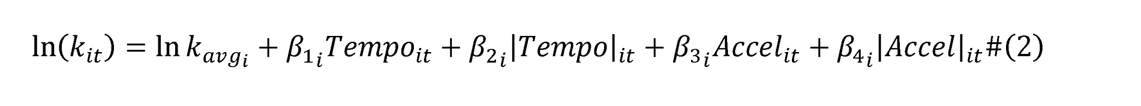

Participants’ log discount rate on each trial was modeled as a sum of that participant’s overall average log discount rate (*lnk*_avgl_), and four task-dependent effects.

The first two terms deal with the effect of time-counting pace on discount rate. The term *r,m,,*_it_ is coded so that fast condition trials are 1, normal condition trials are 0, and slow condition trials are -1, thereby modeling the linear effect of counting tempo on discount rates. Previous research has shown that fast-tempo music led to higher discount rates than slow-tempo music, which would predict discount rates to vary in the order of fast, normal, and slow conditions (K. Kim & Zauberman, 2019). On the other hand, our time-counting task involves effort on the participant’s part to match the tempo, which may lead to higher effort and arousal in fast and slow conditions because people may be more comfortable counting time in 1 second intervals. To account for this effect, we include the absolute value of the term *r,m,,*_it_ so that both the fast and slow conditions are coded as 1, while the normal condition is 0.

The last two terms, *Acc,,*_it_ and *|Acc,,|*_it_, deal with the effect of, not the pace of time-counting itself, but the change in the pace of time-counting. The motivation is that whether a tempo feels ‘fast’ or ‘slow’ may be more relative than absolute, such that counting time every 1 second may feel fast when it comes after counting every 1.3 seconds, but it may feel slow when it comes after counting every 0.7 seconds. To model this temporary effect occurring from transitioning between conditions, the term *Acc,,*_it_ models the first 15 trials after a change in condition, which happens twice in each run. The very first trial after change in condition is given the largest weight at 1 if the pace is accelerating and -1 if the pace is decelerating; the following 14 trials are given a linearly diminishing weight until weight reaches 0 at the 16^th^ trial. The term *|Acc,,|*_it_ takes the absolute value of *Acc,,*_it_ , and hence models any condition transition with positive weights. This regressor is akin to *|r,m,,|*_it_ in that it models any effect of effort or arousal that comes from sudden changes of tempo which may temporarily make the task more difficult. We also considered alternative formulations of the *Acc,,*_it_ regressors and have found that the results are largely the same (**supplemental materials**).

All individual parameters were linked to a group level normal distribution:

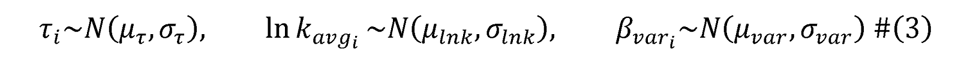

The model was fit using RSTAN with 4 chains with 5000 samples each, 2000 burn-in. No priors were declared for any of the parameters, which meant that they had an improper uniform prior over the real support (or positive support for *σ*_var_ and *T*_i_). All parameters had r-hat of 1 after concluding the sampling, indicating that the chains have sufficiently mixed and converged. Significance testing for each regressor was performed at the group level by examining whether the 95% Bayesian interval for the group mean ( _var_) for each regressor contained zero.

For clarity, we would like to note that not all four regressors are meant to model the manipulation of ‘time perception’. Only *r,m,,*_it_ and *Acc,,*_it_ assume a linear relationship between time-counting pace and discount rate (former being absolute pace and latter being relative pace). The other two regressors, while they may affect discount rates, are likely modeling the effects of other factors such as arousal, difficulty, or surprise which may also simultaneously affect discount rates.

## Image acquisition and preprocessing

The images were collected with a Siemens 3T Trio scanner with a 32-channel head coil. High-resolution T1-weighted anatomical images were acquired using an MPRAGE sequence [repetition time (TR) = 1,900ms; echo time (TE) = 2.52 ms, 256 axial slices, 1.000 mm isotropic voxels; 192 x 256 matrix). T2*-weighted functional images were acquired using an EPI sequence with 42 axial slices 3mm isotropic voxels, 80 x 80 matrix, TR = 2,000 ms, TE = 20 ms, in an interleaved acquisition sequence.

All images were preprocessed using fMRIPrep 20.2.1 (Esteban et al., 2019). EPI sequences were skull-stripped, co-registered using boundary-based registration with nine degrees of freedom, head-motion corrected via six degrees of freedom, slice-time corrected, and normalized to a 2mm MNI space. Before running the GLM model below, the images were smoothed with a FWHM 8mm Gaussian kernel.

## GLM analysis

The second step in our analysis is to test if the behavioral manipulations of time perception also alter activity in brain regions previously implicated in time-keeping. Furthermore, if the manipulation was truly successful in altering time perception, which alters discount rates, we should expect significant neural activity change in both the time-counting task and the ITC task.

We used a GLM to model the fMRI time-courses in both the time-counting task and the ITC task simultaneously (i.e., in one model). This not only allows us to find neural signals that correlate with various time-counting manipulations in both tasks, but also ensures that the signal that we find in one task is not simply a spillover from the other task’s signal that hasn’t been accounted for. Specifically, we investigated the neural correlates of the two time-counting manipulations that significantly affected discount rates: *|r,m,,|*_it_ and *Acc,,*_it_.

For the time-counting task, we modeled the 5 second period leading up to the button press as this is the time at which the internal pace of time-counting is most pertinent to the response (as opposed to earlier periods of the task where participants are learning the pace from external stimuli). We used an overall boxcar event regressor to model the average response in this period and added the two modulator regressors *|r,m,,|*_it_ and *Acc,,*_it_ to model the fluctuations in activity due to the time perception manipulations. For the ITC task, we modeled the 2 second period from the onset of the trial using one overall boxcar event regressor and two modulator regressors. In addition, we also included a subjective value regressor in order to control for regions whose signal is primarily subjective value. While subjective value signals would also be affected by our manipulations, we are more interested in regions that hold information about time perception, not valuation. The trial-by-trial subjective value was calculated as *u*_it_ term in **eq. 1**.

Twelve motion parameters (six affine transforms and their squares) were also included as nuisance regressors to control for motion artifacts. The significance testing for each regressor was performed at the whole-brain level using cluster extent permutation tests with cluster forming threshold of *p* = .005 at false discovery rate (FDR) of *q* = .05. We used the permutation test (as opposed to GRF) to control the FDR exactly even at cluster forming thresholds of p = .005.

## Brain-behavior connection

The final step in our analysis is to show that the degree of neural activity change in time- keeping regions caused by our experimental manipulations are predictive of the degree of change in behavioral discount rates caused by the same manipulations. To do this, we searched for brain regions in which the change in neural activity due to the time perception manipulations predicted the change in log discount rate in behavior due to the same manipulation. We first performed a voxel-level correlation between the individual behavioral coefficients obtained from model in **eq. 2** (i.e., *1*_2i_ , *1*_3i_) and the corresponding individual neural contrasts obtained from the GLM analysis for ITC. Then, we performed a whole-brain correction using cluster extent permutation testing with cluster forming threshold of *p* = .005 at false discovery rate (FDR) of *q* = .05.

## Results

### Behavioral results – time-counting task

Participants responded appropriately to the changing tempo in the time-counting task. The participants’ average counting pace was 0.73 seconds (SD = 0.04) in the fast condition, 1.01 seconds (SD = 0.06) in the normal condition, and 1.25 seconds (SD = 0.07) in the slow condition (**Figure 2A**). These closely matched each condition’s correct pace at 0.7, 1, and 1.3 seconds, respectively. Notably, we found that participants were counting generally slower than 0.7 seconds in the fast condition, while generally faster than 1.3 seconds in the slow condition, suggesting that they may be more comfortable with counting at a 1 second pace. From visual inspection, participants seemed to quickly adapt to the tempo of the condition from the very first trial of each condition (**Figure 2B**).

**Figure 2.**
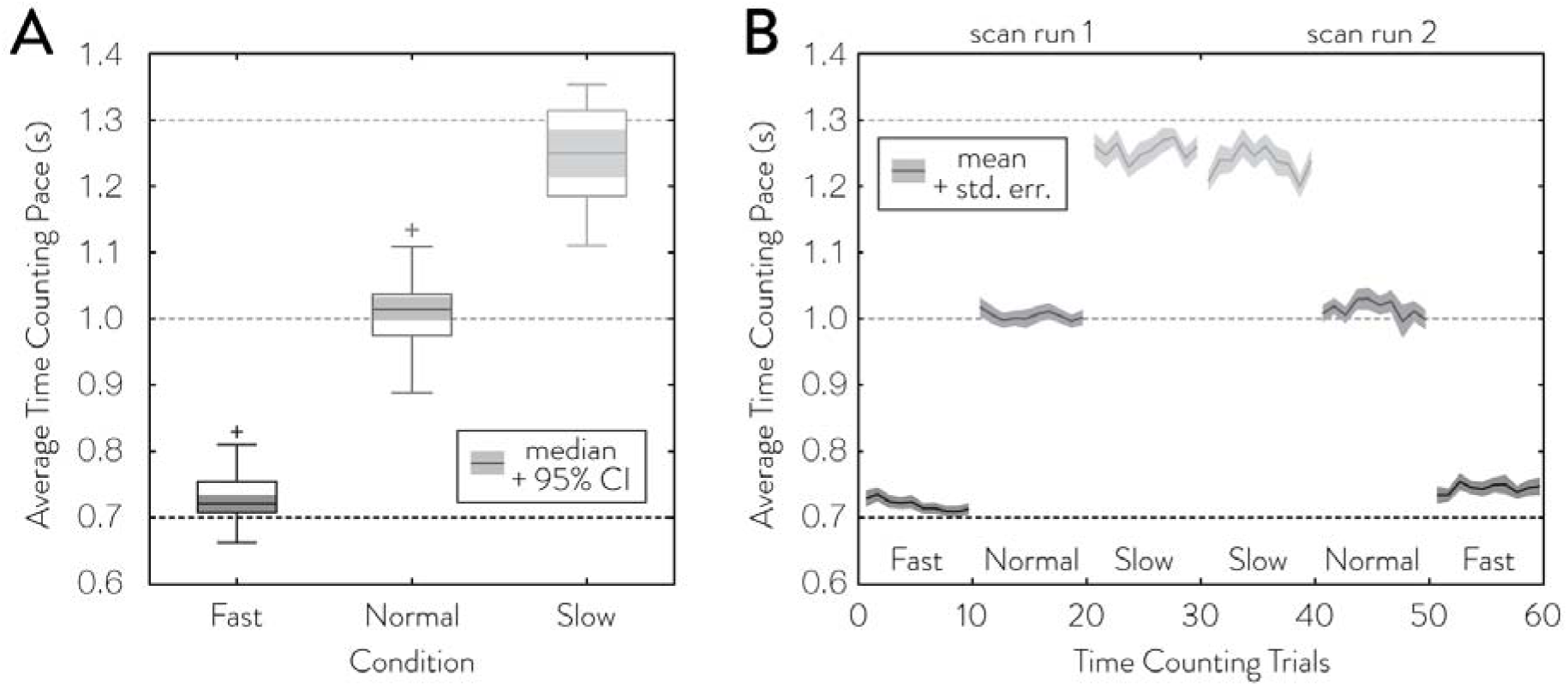
Time-counting task behavioral results. Panel A shows the average response times of time-counting task grouped by the conditions in the time-counting task. The bottom and top of the box denotes the interquartile range, the whiskers denote the full range of data except for the outliers (>3 standard deviations) marked with a cross. The middle line in the box is the median and the shaded area around it is its 95% confidence interval. Panel B shows the average response times of time-counting task for each trial. The solid line denotes the mean while the shaded area denotes the standard error of the mean.

### Behavioral results – ITC task

To identify the time-counting manipulation that alters discount rates, we modeled the participants’ discount rates over time as a function of the time-counting conditions and the change between the conditions.

Our behavioral model provided a good fit to the participants’ choices in the ITC tasks. We assessed the model’s in-sample accuracy using the area under the receiver operating characteristic curve (AUROC), which serves as a threshold-free accuracy. We used AUROC instead of raw accuracy as it maintains 50% as the at-chance null accuracy even when the choices are not equally balanced (e.g., for someone who is heavily impatient, blindly predicting a one-sided choice for all trials will lead to a raw accuracy that is over 50%). On average, our model was able to predict 98% of the choices that the participants made (range 92.9% - 99.7%), and our pseudo-R^2^ metric was high for each subject’s data with an average of 0.79 (range 0.555 – 0.903). The participants’ average discount rates were distributed within the range of -7 and -2, which was well within the range of discount rates (-7.5 and 1) our task choices were designed to measure.

The first step in our analysis is to show that our experimental manipulations of the participants’ time-counting pace significantly alters their discount rates. We investigated four different ways in which the discount rate may be affected: linear effect of tempo (counting pace), absolute effect of tempo, linear effect of tempo acceleration, and absolute effect of tempo acceleration. **Figure 3A** graphically shows the four regressors used to model the discount rates as a function of these four effects. Of the four tested, we found two task effects that significantly affected the participants’ discount rate at the group-level. Firstly, participants’ discount rates were significantly higher in the fast and slow conditions compared to in the normal condition (**Table 1**, _|tempo|_ mean = 0.17, Bayesian 95% interval: [0.07, 0.27]). On average, participants had the highest log discount rates in the slow condition (+ 0.21 from normal), followed by fast (+0.13 from normal) and then normal conditions (**Figure 3B**). Congruently, the group-level estimate was not significant for the linear effect of time counting pace on discount rates (**Table 1**, _tempo_ mean = -0.04, Bayesian 95% interval: [-0.10, 0.02]). Secondly, participants’ discount rates were significantly affected by pace acceleration such that the discount rates had a phasic decrease when transitioning to a slower condition and a phasic increase when transitioning to a faster condition (**Table 1**, _accel_ mean = 0.2, Bayesian 95% interval: [0.03 0.35]; **Figure 3B**). The absolute effect of acceleration was not significant at the group level (**Table 1**, _|accel|_ mean = 0.09, Bayesian 95% interval: [-0.06 0.24]).

**Figure 3.**
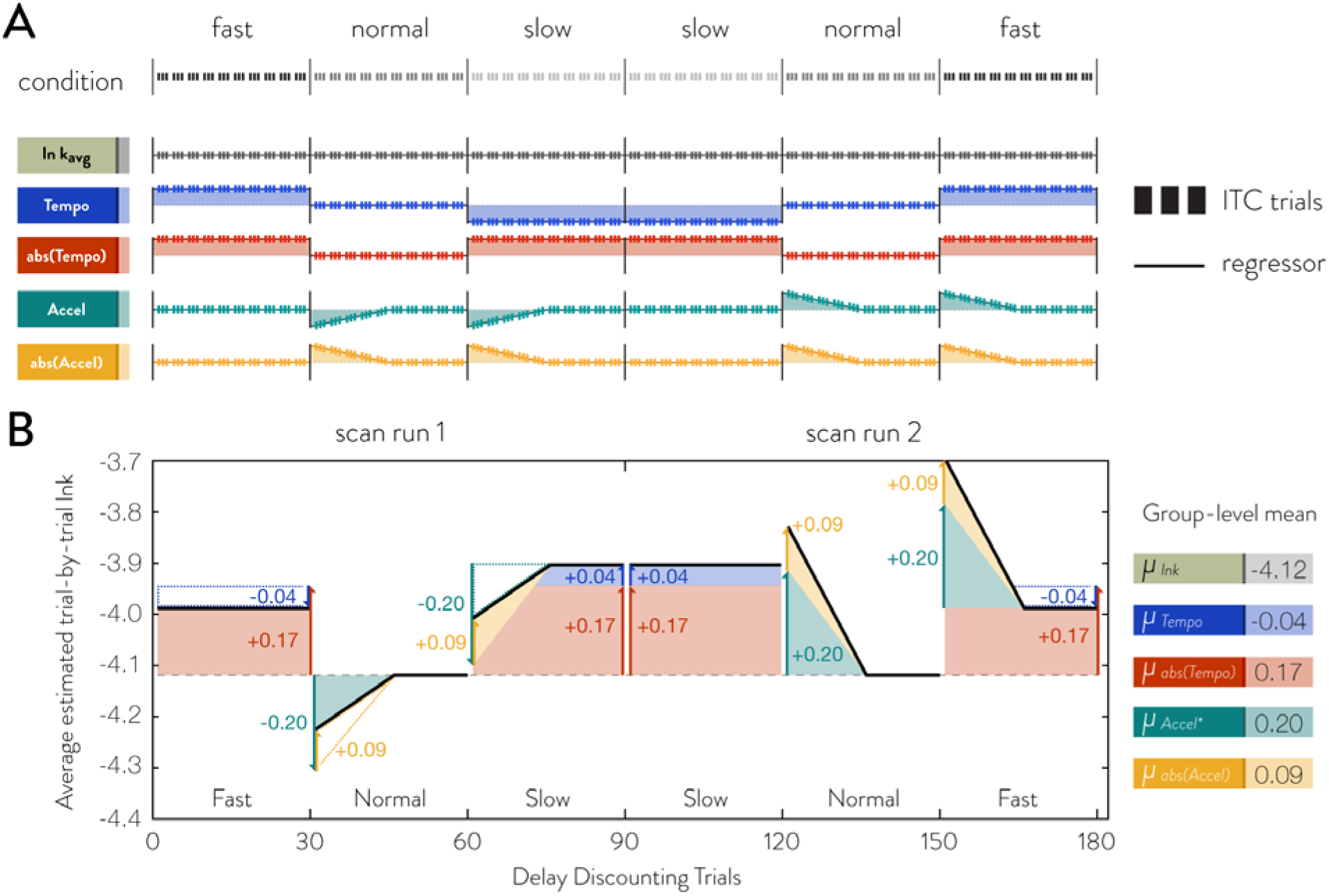
Delay discounting task – effects of time-counting manipulation on trial-by-trial discount rates. Panel A shows the time-counting manipulation regressors used to model the trial-by-trial discount rate in the ITC task. The vertical triplet bars indicate the three delay discounting trials of a block, while the solid lines through the bars depict the regressor level. Panel B visually illustrates the estimated group average trial-by-trial log discount rates in black solid line with the additive effects of the regressors marked in shaded areas and arrows. The dotted grey line denotes the group average log discount rate.

**Table 1.**
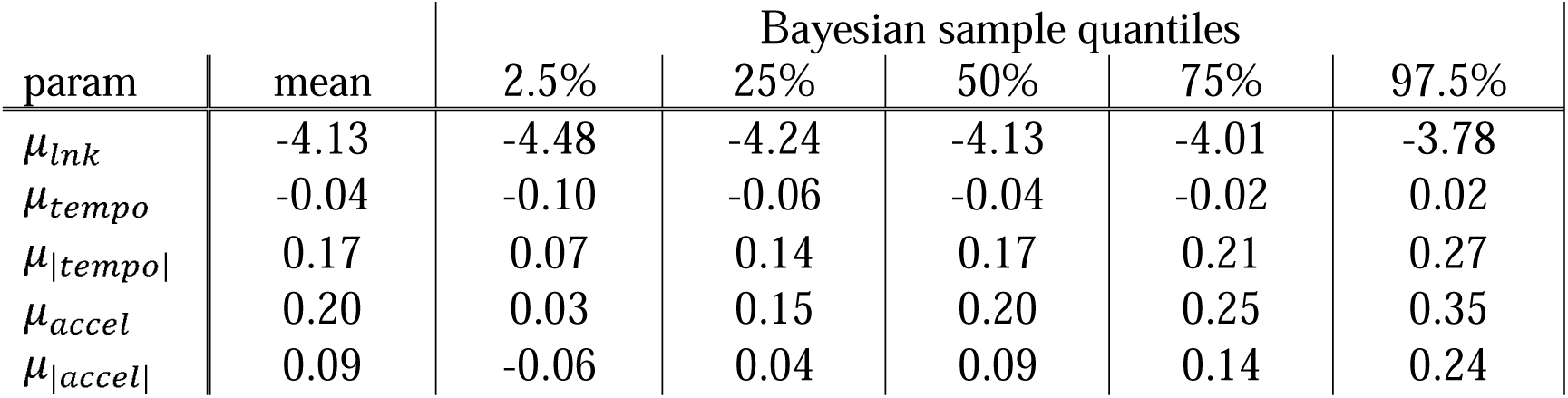
Bayesian hierarchical model group mean estimates. Posterior sample means and quantiles are shown for group level parameters from equation 3. From above to below are group average log discount rate, followed by five regressors corresponding to each of the five hypotheses.

### Imaging results

In the time-counting task, we found no brain regions whose activity was significantly different between the normal conditions versus the fast and slow conditions (**Fig. 4A**). However, a number of brain regions were shown to be negatively related to phasic effects of pace acceleration (**Fig. 4B**). These regions included the left anterior insula, dorsomedial prefrontal cortex (dmPFC), precuneus, and bilateral superior temporal gyri. Activity in these regions temporarily increased after switching to a slower time-counting condition and temporarily decreased after switching to a faster time-counting condition.

**Figure 4.**
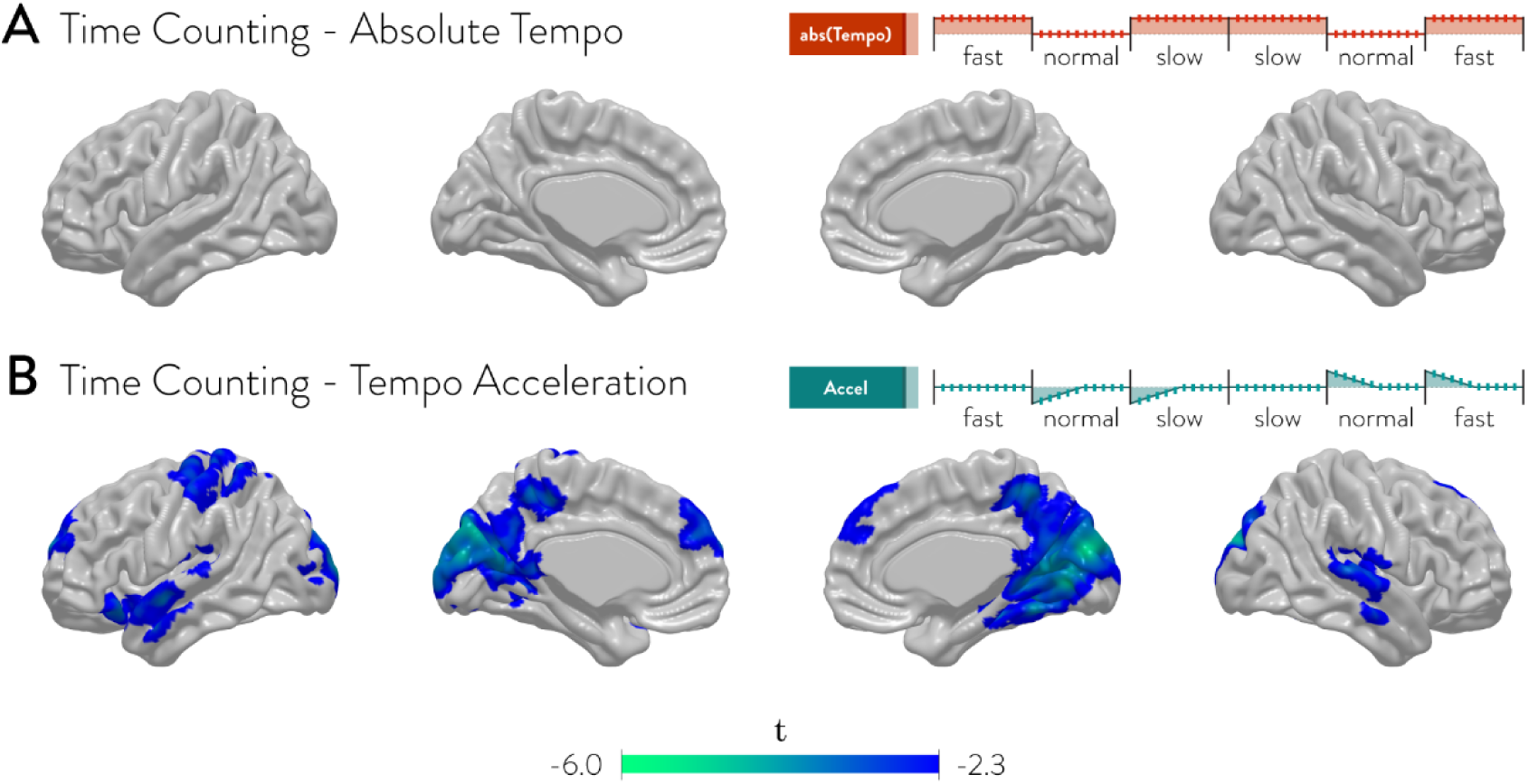
fMRI GLM results – Time-counting task. Shown above are brain regions that have significant activity correlated to each of the two regressors in time-counting task. All results were permutation tested at the whole-brain level.

A similar pattern was also found in the ITC task. No brain regions were significant for the contrast between the normal conditions versus the fast and slow conditions (**Fig. 5A**). Again, however, we found a diverse set of brain regions that showed activity significantly correlated with phasic effects of time-counting pace acceleration (**Fig. 5B**). There was extensive overlap with the brain regions showing a similar effect in the time-counting task, including the left anterior insula, dmPFC, precuneus, and bilateral superior temporal gyri. These regions were negatively related to the pace acceleration such that their activities increased when transitioning to a slower time-counting condition and decreased when transitioning to a faster time-counting condition.

**Figure 5.**
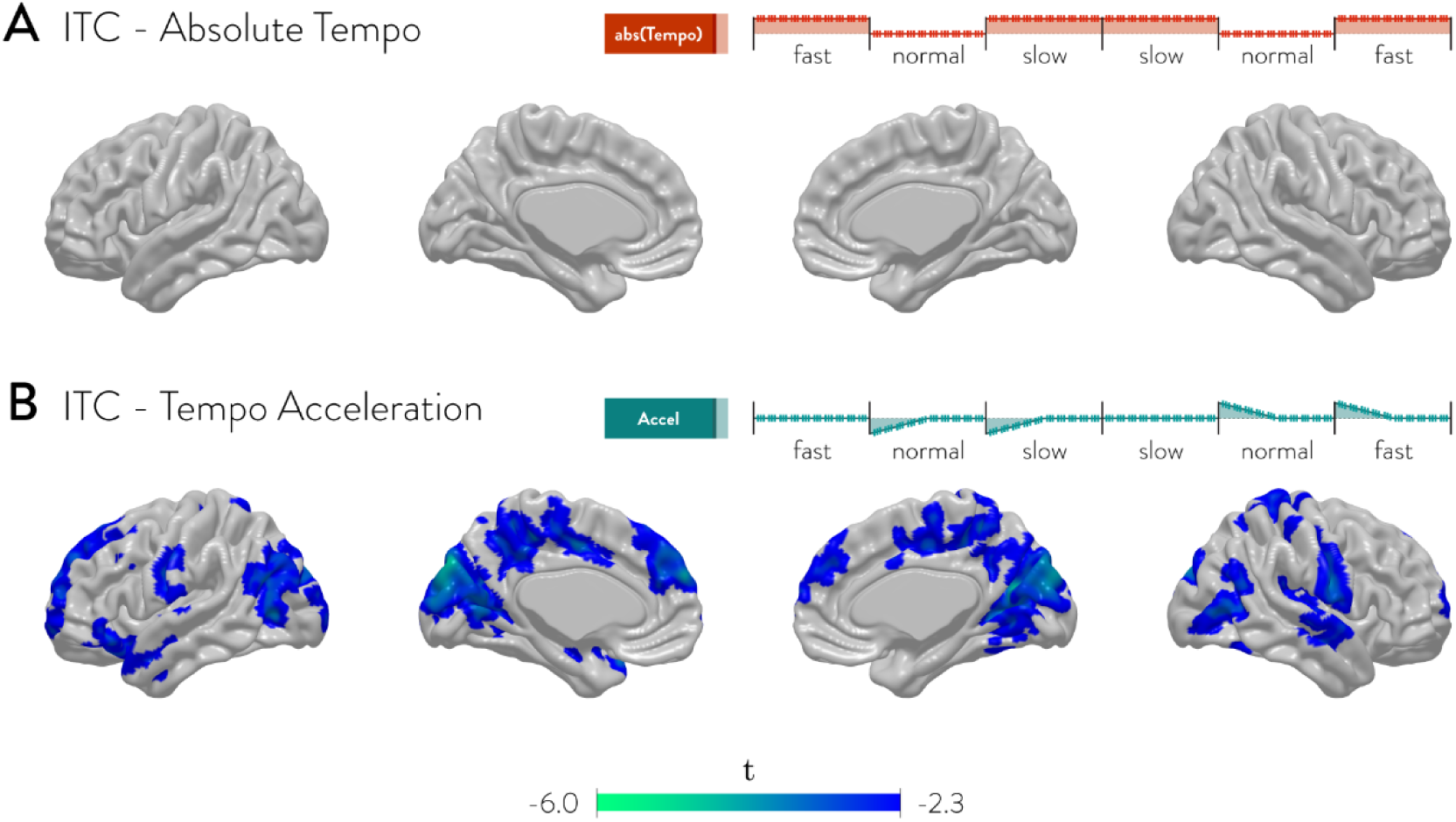
fMRI GLM results – ITC task. Shown above are brain regions that have significant activity correlated to each of the two regressors in ITC task. All results were permutation tested at the whole-brain level.

### Brain-behavior connection

We found four brain regions, at the whole-brain correction level, in which greater neural changes in response to the time-counting manipulation predicted greater behavioral changes in discount rate (**Fig. 6**). These regions included the left anterior insula, the right temporal gyri (superior, middle, inferior), the right occipital pole and the left middle occipital lobe. In each of these four regions, the effect of condition change on neural activity was negatively correlated with the effect of condition change on log discount rate. This meant that people who showed greater decrease and increase of discount rates following a slowing and quickening condition change (respectively) also showed greater increase and decrease (respectively) of neural activity in these regions. More strikingly, the linear relationship between the behavioral and the neural measures closely passed the origin point in all four regions such that a negative behavioral effect corresponded to a positive neural effect while a positive behavioral effect corresponded to a negative neural effect; this result goes beyond a simple correlative relationship as it means that the sign of the change in neural activity is also predictive of the direction of change in log discount rates.

**Figure 6.**
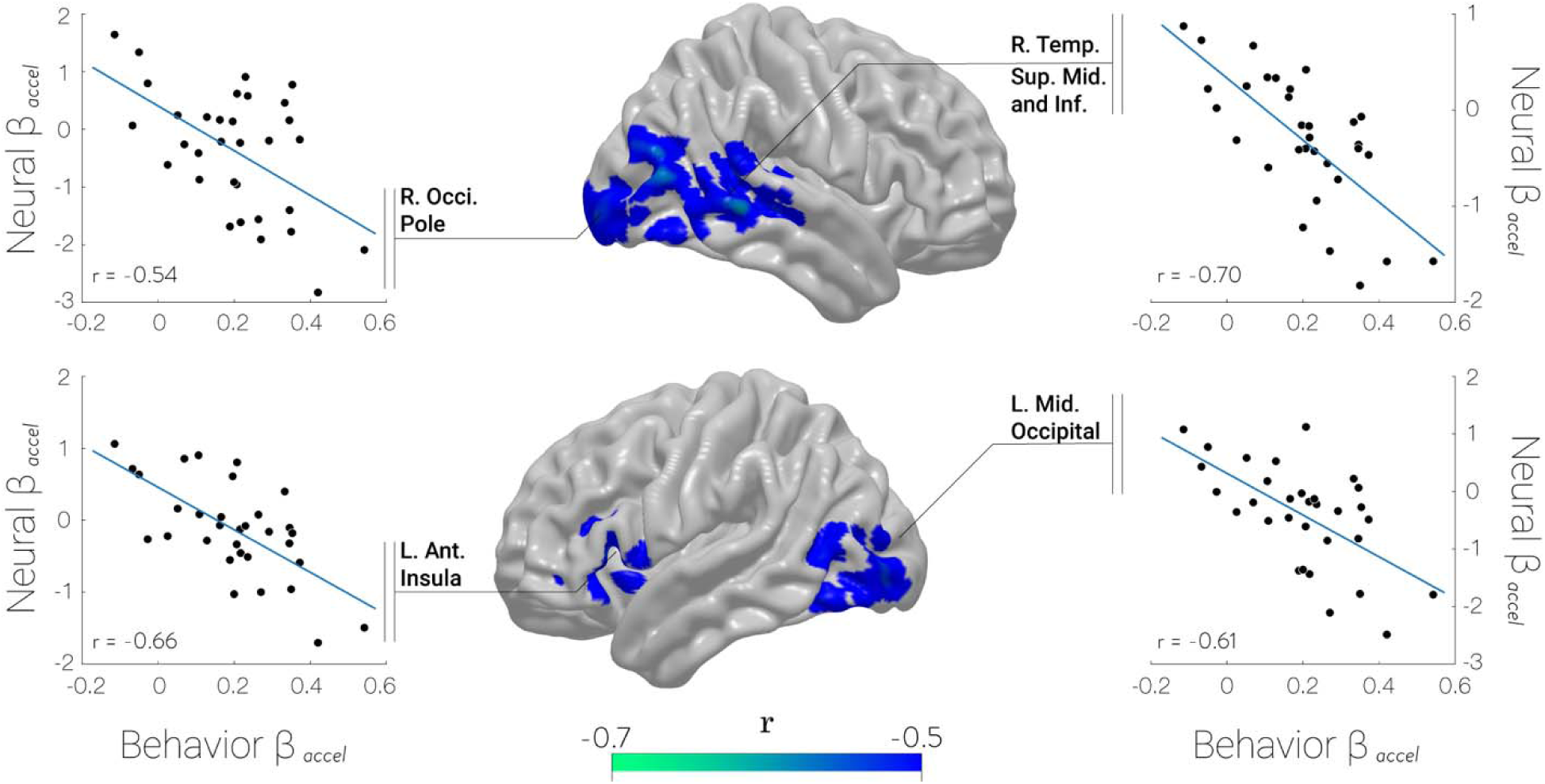
Link between brain and behavior – correlation between effects of time-counting pace acceleration on individual fMRI activity (ordinate) and on individual log discount rate (abscissa) during ITC. Shown above are brain regions that have significant correlation at the whole-brain level between neural response to time-counting pace acceleration (i.e., Fig. 5B), and change in log discount rate due to time-counting pace acceleration (i.e., Fig. 3A _accel_). Regions were permutation tested at whole-brain level and hence p-value in each of the respective scatterplot is omitted as regions have already been selected based on a whole-brain test (but they are all equal to *p* = .001 or below).

## Discussion

In the current paper, we investigated the neural underpinnings of how our internal time perception influences our discount rates by manipulating the former and observing the effect on the latter. We followed a three-step procedure to 1) show that a time-counting manipulation changes discount rates, 2) show that the same manipulation also affects neural activity, and 3) show that the degree of neural activity change due to manipulation predicts the degree of discount rate change due to that same manipulation.

Starting with the first of our main analyses, we found that when participants switch time-counting conditions from a faster counting one to a slower counting one, their discount rates temporarily decreased; whereas switching conditions from a slower counting one to a faster counting one led to a temporarily increased discount rate. While we also found that the speed of counting itself also affected discount rates, the participants’ discount rates were increased in both fast and slow counting conditions compared to baseline, showing that the speed of counting time itself is not linearly related to changes in discount rates. These results suggest that the relative transition between different speeds of counting time leads to changes in the participants’ time perception rather than the rote speed of counting time itself. It is difficult to exactly isolate why peoples’ discount rates were increased in both fast and slow conditions, especially given the lack of any corresponding brain activity changes, but it’s possible that factors such as arousal from task difficulty may play a role as participants were generally less accurate for counting at fast (0.7 sec) or slow (1.3 sec) speeds compared to normal speed (1 sec).

Having shown that time-counting pace acceleration affects discount rates, we then assessed if that manipulation also affected neural activity in both the time-counting and ITC tasks in the time-related regions. We found that the neural activity in left anterior insula, dmPFC, precuneus, and bilateral superior temporal gyri, for both time-counting task and ITC task, temporarily increased after switching to a slower counting condition and temporarily decreased after switching to a faster counting condition. These results also add further support that our experimental manipulation has affected the participants’ internal time perception as evidenced by corresponding activity change in insula and superior temporal gyri, both of which has been previously found to show activity corresponding to an internal clock (Bueti & Macaluso, 2011; Wencil et al., 2010; Wittmann et al., 2010, 2011).

Finally, we found brain-behavior correspondence where both the direction and the degree of change in neural activity in insula and temporal gyri following a condition switch was predictive of the direction and degree of change in log discount rate following a condition switch. Especially given that the anterior insula has been theorized as a crucial locus of perceiving time (Craig, 2009a, 2009b), future studies may employ neural manipulation designs such as TMS (transcranial magnetic stimulation) or tDCS (transcranial direct current stimulation) to test if activity changes in these regions alone can alter discount rates.

We would like to emphasize that we do not believe time perception is the sole process in determining people’s discount rates. In fact, even within our findings, we found that the rote speed at which people count time affects peoples’ discount rates but not in a way that time perception would affect them. Delay discounting is likely a multifaceted process where there are numerous factors that can influence discount rates at various points of the decision process (Addis et al., 2011; Lee et al., 2022; Liberman & Trope, 2014; Palombo et al., 2015; Peters & Büchel, 2010; Rösch et al., 2022; Soman et al., 2005; Trope & Liberman, 2010; Urminsky, 2017).

As a newly developed task, there are several aspects of our experimental design that we believe will benefit from modifications in future research. In current research, the change in time-counting speed was only between fast-normal and normal-slow conditions but not fast-slow conditions, as we feared it may be too disorienting or jarring for participants to make a larger change in speed. Nevertheless, there is a standing question of whether a greater change in time-counting speed leads to a commensurately greater change in discount rates. Furthermore, because the current design required a relatively large number of trials in each condition (30 trials) in order to distinguish the condition effects and phasic effects, it limited the number of testable conditions and transitions between conditions. Future research may benefit from intertemporal bidding paradigms as this would allow for a more direct observation of a trial-by-trial discount rate without relying on modeling.

## Acknowledgements

This work was supported by the National Research Foundation of Korea (NRF) grant funded by the Korean government (MSIT) (NRF-2022R1A2C1010704). This funding source played no role in the study design, the collection of data, the analysis and interpretation of the data, the writing of the manuscript, and the decision to submit the article for publication. The authors thank Jason (Sangyoon) Jung, Stella (Sanghyo) Jung, and Ji Yeon Han for their continued encouragement.

