## Supplemental Materials for "Altering subjective time perception leads to correlated changes in neural activity and delay discounting impulsivity"

### *Behavioral Data Analysis – Alternative model*

To check the robustness of our results, especially for $Accel_{it}$ and $|Accel|_{it}$, we also considered an alternative model where the transient effect of time-counting tempo change is modeled only for the first 3 blocks after condition change (as opposed to a linearly decreasing effect as in the main manuscript). Here, the first 9 trials after condition change is given the weight of -1 after a decelerating transition and the weight of 1 after an accelerating transition (**Figure S1**).

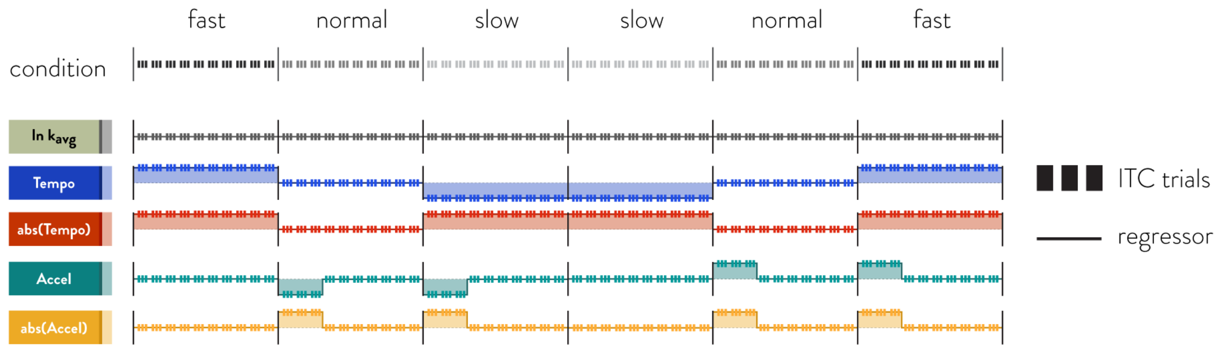


**Figure S1. Delay discounting task – alternative time-counting manipulation regressors.** Panel A shows the time-counting manipulation regressors used to model the trial-by-trial discount rate in the ITC task. The vertical triplet bars indicate the three delay discounting trials of a block, while the solid lines through the bars depict the regressor level.

Using the same hierarchical Bayesian framework as in the main manuscript, we found the overall pattern to be similar as in the main manuscript (**Table S1**). The overall changes in the coefficients were very minor and it was still the case that only the two regressors, $|Tempo|$ and $Accel$, were significant at the group level.

| | | Bayesian sample quantiles | | | | |
|---|---|---|---|---|---|---|
| param | mean | 2.5% | 25% | 50% | 75% | 97.5% |
| $\mu_{lnk}$ | -4.12 | -4.48 | -4.24 | -4.12 | -4.00 | -3.77 |
| $\mu_{tempo}$ | -0.04 | -0.09 | -0.06 | -0.04 | -0.02 | 0.02 |
| $\mu_{\|tempo\|}$ | 0.17 | 0.07 | 0.14 | 0.17 | 0.21 | 0.27 |
| $\mu_{accel}$ | 0.19 | 0.05 | 0.14 | 0.19 | 0.23 | 0.32 |
| $\mu_{\|accel\|}$ | 0.06 | -0.07 | 0.01 | 0.06 | 0.10 | 0.18 |

**Table S1. Bayesian hierarchical model group mean estimates.**
